# Structural insight into bacterial co-transcriptional translation initiation

**DOI:** 10.1101/2024.03.19.585385

**Authors:** Takeshi Yokoyama, Yuko Murayama, Tomomi Uchikubo-Kamo, Yuri Tomabechi, Asuteka Nagao, Tsutomu Suzuki, Mikako Shirouzu, Shun-ichi Sekine

## Abstract

In bacteria, transcription and translation are tightly coupled, forming a transcription-translation complex (TTC) between RNA polymerase (RNAP) and the ribosome. As nascent mRNA emerging from RNAP is susceptible to ribonuclease digestion, undesired RNA folding, and R-loop formation, immediate TTC formation is important. Here, we report the cryo-electron microscopy structures that capture the translation initiation complex assembly on transcribing RNAP. As a short mRNA emerging from RNAP, the 30S ribosomal subunit binds the RNAP on its inter-subunit side, interacting with the mRNA 5’-region and the initiator tRNA. The RNAP could relocate to the canonical mRNA-entry site of 30S, threading the mRNA into a path formed between RNAP and 30S. The subsequent 50S joining establishes a TTC. These structures illustrate the transcription-coupled translation initiation, while protecting mRNA.

**One-Sentence Summary:** Cryo-EM captures the ribosome assembly on transcribing RNA polymerase in the presence of translation initiation factors.

## Introduction

Gene expression is comprised of two fundamental processes in all organisms: transcription and translation. Genetic information encoded as a nucleotide sequence on DNA is transcribed into mRNA by RNA polymerase (RNAP), and subsequently translated into its corresponding amino-acid sequence by the ribosome (*1*)(*2*). In eukaryotes, these two processes occur independently in different compartments of the cell, separated by the nuclear membrane (*3*). By contrast, in prokaryotes, both two principal gene-expression processes occur in the same time and space, and they are tightly coupled (*4*)(*5*). Early electron microscopic studies visualized that ribosomes are attached to mRNAs that are being transcribed by RNAP migrating on DNA fibers (*4*)(*6*). More recently, several cryo-electron microscopy (cryo-EM) structures unveiled molecular details of the transcription-translation complexes (TTC), visualizing how the trailing ribosome interacts with the leading RNAP (*7*)(*8*)(*9*)(*10*)(*11*). Functionally, there are significant benefits in the transcription-translation coupling. Prokaryotic RNAP synthesizes nascent RNA at the highest speed of 20 - 80 nt s^-1^, with punctuated pauses allowing the co-transcriptional RNA folding (*12*)(*13*). The RNA hairpin formed within the RNAP exit channel triggers an elemental pause by keeping RNAP inactive in a swiveled (or ratcheted) conformation (*14*)(*15*)(*16*)(*17*). Upon coupling, the leading ribosome exerts a forward-directing force on the RNAP, which allosterically induces RNAP into an anti-swiveled conformation (*18*). This facilitates efficient RNAP translocation with less frequent pausing during transcription, avoiding the situation of ribosome-ribosome collision, which is recognized by the SmrB endonuclease for the ribosome rescue pathway (*19*).

Despite the accumulation of knowledge about translation elongation in TTC, the assembly process of TTC, in which translation initiation occurs on transcribing RNAP, has remained elusive. mRNA from the majority of bacterial genes is supposed to be immediately incorporated into the ribosome 30S subunit as the ribosome binding site (RBS) emerges from the RNA-exit channel of RNAP (*5*)(*7*). There are potential benefits for this rapid incorporation: protection against nascent mRNA degradation by ribonucleases (*20*)(*21*)(*22*) (*23*)(*24*), preventing undesired RNA folding with nearby segments (*13*), preventing mRNA from annealing to the template DNA, forming R-loops (*25*), and preventing premature transcription termination by the transcription termination factor Rho (*26*)(*27*). A nascent elongating transcripts sequencing (NET-seq) experiment showed that RNAP frequently pauses around the translation start sites (*28*), and the RNAP pausing would provide a window of opportunity for the establishment of the transcription-translation coupling. However, it remains unclear how nascent mRNA emerging from RNAP is incorporated into the 30S ribosomal subunit to initiate translation. In this study, we employed cryo-EM to analyze detailed molecular interplay during translation initiation complex assembly on transcribing RNAP. The molecular details of sequential mRNA incorporation process into the 30S/70S ribosome illuminate how TTC is established to initiate co-transcriptional translation.

## Results

### The 30S ribosomal initiation complex formation on RNAP-EC

To investigate how nascent mRNA emerging from RNAP is incorporated into the 30S ribosomal subunit during the 30S initiation complex formation, we performed cryo-EM structural analysis of the RNAP-30S complex in the presence of a set of the translation initiation factors and the initiator tRNA (*29*). In this complex, a 43 nt mRNA was employed. The 5’-end of the mRNA contains RBS with the Shine-Dalgarno (SD) sequence and the AUG initiation codon separated by a 6-nt spacer, while the 3’-end contains a 10-nt sequence to form a 10-bp DNA/RNA hybrid within the RNAP active site (fig. S1). We first reconstituted an elongation complex of RNAP (RNAP-EC) using RNAP from the bacterium *Thermus thermophilus*. Then, the RNAP-30S complex was assembled by incubating the RNAP-EC with the *T. thermophilus* 30S ribosomal subunit, translation initiation factors IF1, IF2, IF3, and fMet-tRNA^fMet^ (Fig. 1A). The reconstituted complex was examined by single particle analysis (Fig. 1 and fig. S2). After extensive image classification, we obtained several 30S complexes having the RNAP densities located at the inter-subunit surface of the 30S ribosome with various combinations of the translation initiation factors (fig. S2A). One of these structures yielded a clear cryo-EM density for RNAP, which is stably located at an unprecedented position straddling the 30S shoulder and beak (Fig. 1B). Henceforth, we will refer to this complex as “complex 1” for simplicity. In complex 1, mRNA emerging from the RNAP is accommodated into the mRNA path on the 30S subunit, forming SD-antiSD duplex and fMet-tRNA^fMet^ binding at the P site (*30*)(*31*)(*32*)(*33*). While the 5’-end of mRNA, from the SD sequence to the second UUU codon at A site, is accommodated in the mRNA path of the 30S subunit, the following portion of the mRNA comes out from the path to reach the RNAP (Fig. 1, C, D and H). In this complex, the RNAP ω subunit is located in close proximity to IF1 bound to the 30S (Fig. 1B), while the RNAP β’ subunit contacts helix 33 of the 16S rRNA located at the beak (fig. S3A). This unprecedented RNAP-30S binding mode in complex 1 represents how the nascent mRNA is captured by the 30S subunits immediately after it emerges from the RNAP exit channel.

**Fig. 1.**
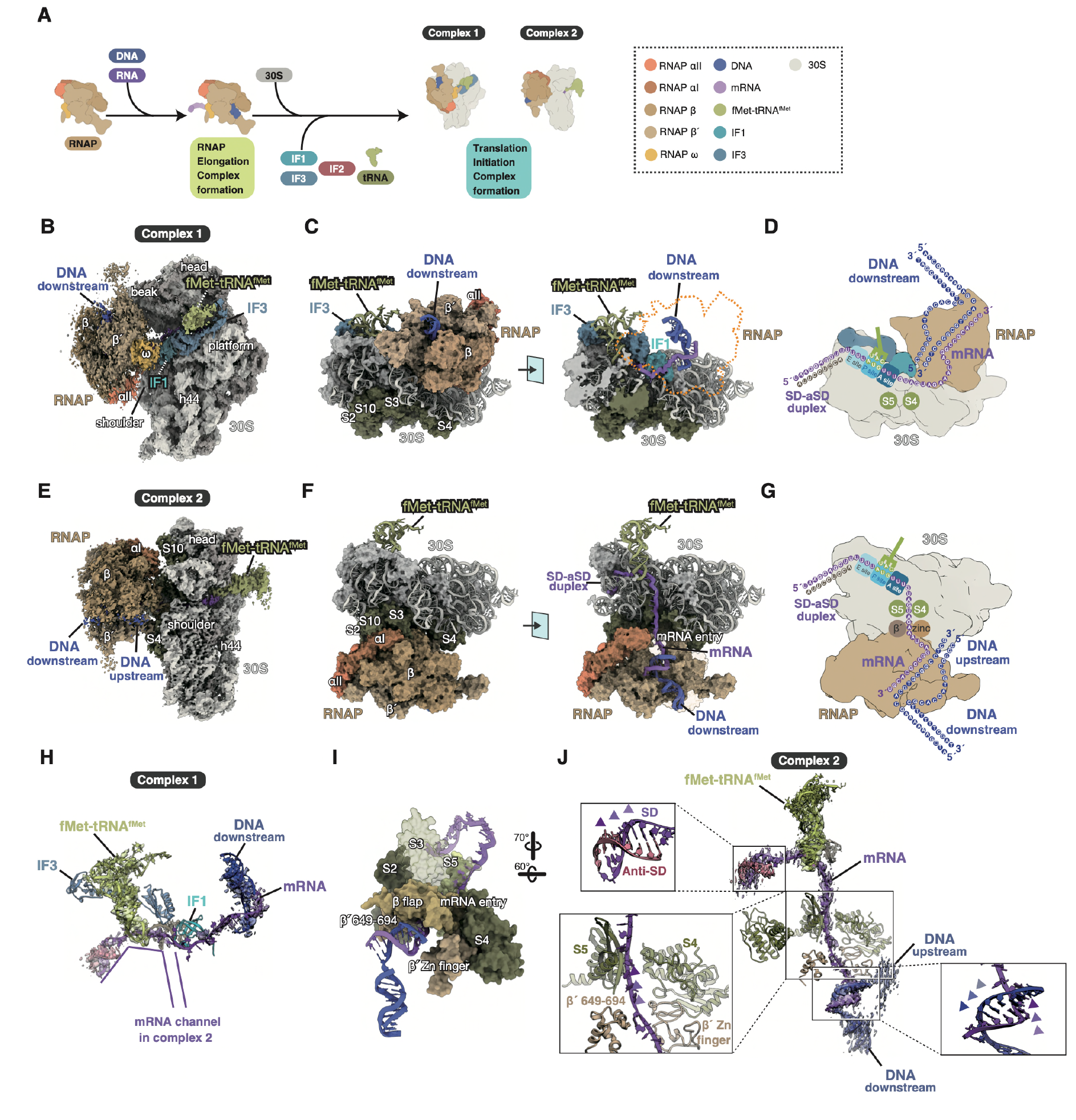
Cryo-EM structures of the 30S initiation complex assembled on transcribing RNAP. (**A**) Schematic representation of the assembly of the 30S initiation complex on RNAP-EC in the presence of translational initiation factors for cryo-EM analysis. (**B**) Cryo-EM reconstruction of the RNAP-EC bound to the inter-subunit side of the 30S subunit (complex 1). (**C**) Surface representation of the model of complex 1. The right panel shows the cross section of the left structure, showing IF1 and the mRNA path. (**D**) Schematic representation of complex 1 showing the DNA/RNA interaction and location inside the complex. (**E**) Cryo-EM reconstruction of the RNAP-EC bound to the back side of the 30S subunit (complex 2). (**F**) Surface representation of the model of complex 2. The right panel shows the cross section of the left structure, showing the mRNA path. Structures in (**C**) and (**F**) are aligned by 30S as a guide for comparison. (**G**) Schematic representation of complex 2 showing the DNA/RNA interaction and location inside the complex. (**H**) The mRNA path in complex 1 showing interactions with IF1, IF3, and fMet-tRNA^fMet^. (**I**) The surface representation of the mRNA channel with mRNA and DNA in complex 2. (**J**) mRNA in complex 2 showing the SD duplex formation and interaction with the mRNA channel formed by RNAP and the 30S subunit.

In this dataset, we also obtained another complex class (“complex 2”), having the RNAP density on the opposite side of the 30S subunit, as compared with complex 1 (Fig. 1E). In complex 2, the manner of RNAP-binding to the 30S subunit is similar to that in the previously reported TTC structures (collided state), although complex 2 does not contain the 50S subunit (Fig. 1E). RNAP forms direct contacts with ribosomal proteins S2, S3, S4, S5, and S10 (NusE) (Fig. 1, F, G, I and fig. S3B)(*34*), forming a continuous mRNA channel connecting the RNAP-exit path to the 30S ribosome mRNA path (Fig. 1F). The docking interface is formed between the β’ 649-694 region, β’ Zn finger, and β-flap of RNAP and S2, S3, S4, and S5 of the 30S subunit (Fig. 1I). mRNA is accommodated in the channel in an extended conformation, and is perfectly covered within the RNAP-30S complex (Fig. 1J). Positively-charged residues within the mRNA path could support a smooth transfer of the unstructured mRNA from the RNA exit of RNAP to the 30S mRNA entry (fig. S4). The basic amino-acid residues in S3 and S4, which interact with the phosphate groups of the mRNA, are reportedly involved in the mRNA helicase activity of the ribosome (*35*)(*36*). Thus, the direct RNAP-30S interactions in both complexes 1 and 2 would protect mRNA from ribonuclease digestion or intra-/inter-molecular RNA structure formation.

### Transition of the RNAP-EC binding modes relative to the 30S subunit

Structural comparison between complexes 1 and 2 illustrate the transition between these complexes (Fig. 2 and fig. S5). Comparison of the cryo-EM maps of complexes 1 and 2 show a drastic RNAP positional change and its accompanied motion of the 30S head (fig. S5A). The positions of RNAP-EC in the two complexes differ by about 180 degrees, with respect to the 30S subunit (Fig. 2A). This observation suggests that RNAP-EC can be repositioned on the surface of the 30S subunit through a drastic swinging motion from the front side (30S inter-subunit side) to the back side. Focusing on mRNAs, the fulcrum of swing is located at the ribosomal A site, because mRNA is tightly clipped at the P site by the fMet-tRNA^fMet^ binding through the start codon recognition (Fig. 2B). Thus, mRNA could serve as a tether for the transition between the two complexes.

**Fig. 2.**
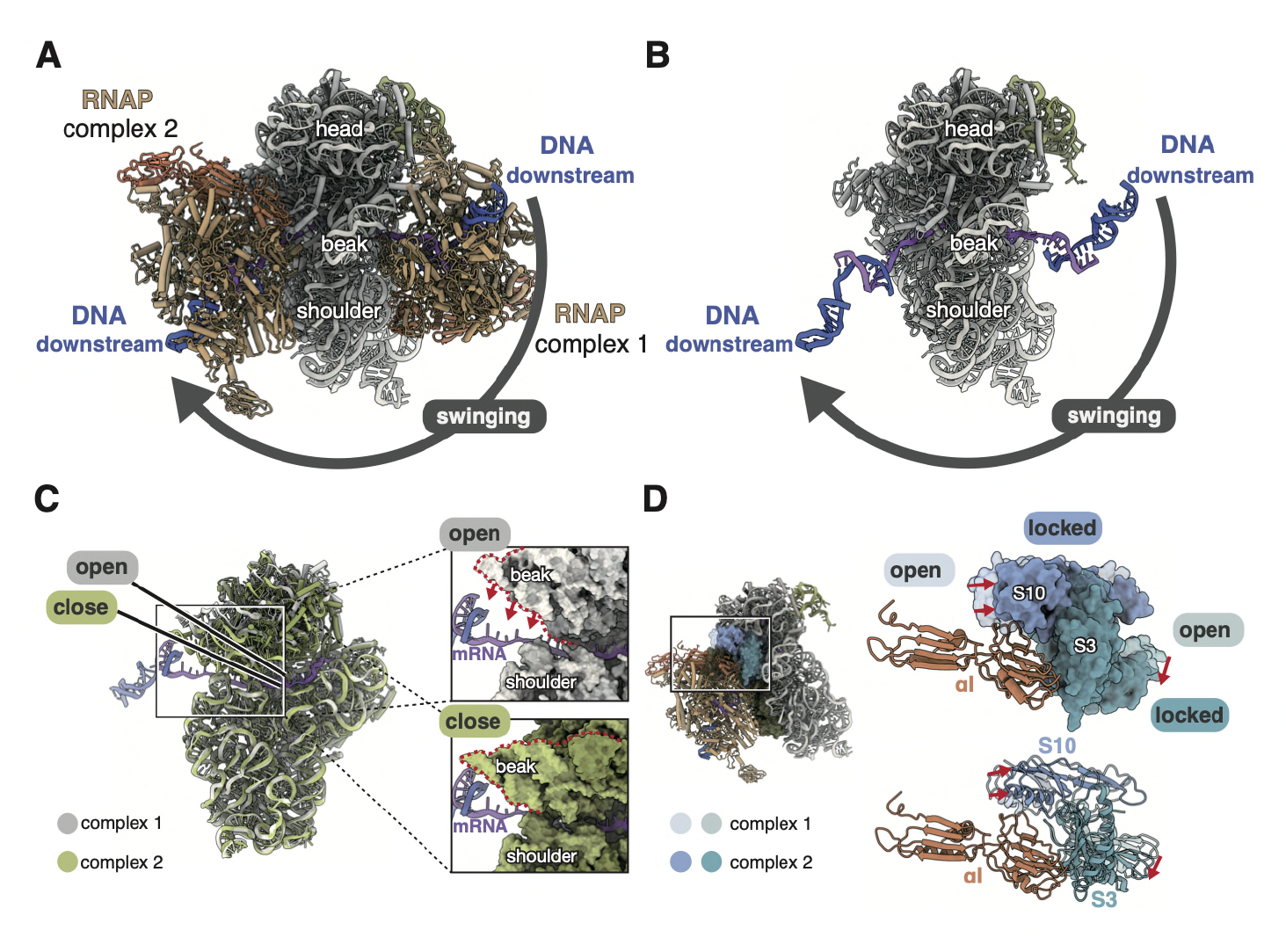
Comparison between complex 1 and complex 2. (**A**) The structural comparison between complexes 1 and 2 suggests that they are switched by swinging of RNAP-EC. (**B**) The same structures without RNAPs showing the mRNA-DNA positions. (**C**) The open and closed conformations of mRNA paths in complex 1 and complex 2, respectively. (**D**) Structural comparison of the ribosomal proteins S3 and S10 between complexes 1 and 2. The RNAP αI subunit forms contacts with S3 and S10 in complex 2.

The head of the 30S subunit could move relative to the body (*37*). This movement changes the space between the beak and the shoulder, which form the mRNA channel between them. In complex 1, this channel is widely open, allowing mRNA to pass through the channel to switch from complex 1 to complex 2 (Fig. 2C). In contrast, in complex 2, this channel is closed due to the shift of the beak. mRNA is held within the closed channel, which seemingly prevents reversion to complex 1 (Fig. 2C). In complex 2, the ribosomal proteins S10 and S3 form contacts with the α subunit (αI) of RNAP. When the 30S head assumed the open conformation, S10 and S3 would cause steric clashes with the RNAP α subunit, suggesting that complex 2 prefers the closed conformation (Fig. 2D). Therefore, it is suggested that the swinging motion of RNAP between complex 1 and complex 2 is ultimately “locked” in complex 2.

### The 70S translation initiation complex assembly on RNAP-EC

Next, we investigated the late step of translation initiation, which establishes a translation elongation competent TTC (*38*). For the reconstitution of TTC, the same 43 nt mRNA used for the 30S-RNAP complex assembly was used (fig. S6). After the RNAP-EC formation, translation initiation complex was assembled including the 50S ribosomal subunit (Fig. 3A). The reconstituted complexes were subjected to the single-particle cryo-EM. After the image processing, we obtained the 70S initiation complex bound with RNAP-EC (“complex 3”) (Fig. 3 and fig. S7). Structural comparison with previously reported TTCs shows that the RNAP-ribosome configuration in complex 3 is similar to that in the collided state (fig. S8)(*10*). In complex 3, the density of IF2 was clearly observed; its domain IV recognizes the CCA end of fMet-tRNA^fMet^ (Fig. 3, B and C). Nearly 70% of tRNA used in this study is charged with formylated methionine, which makes a stable interaction with domain IV of IF2 (fig. S9). Similar to that in complex 2, the mRNA portion embedded in the ribosomal and RNAP environment was clearly observed (Fig. 3, C, D, and E). The extended mRNA connects between the RNAP active site and the SD-anti-SD interaction site within the 30S (Fig. 3E). The interface between RNAP and the 30S subunit is the same as that in complex 2 (Fig. 1I and 3F). This observation suggests that the RNAP-30S interaction is maintained during the 50S incorporation into complex 2.

**Fig. 3.**
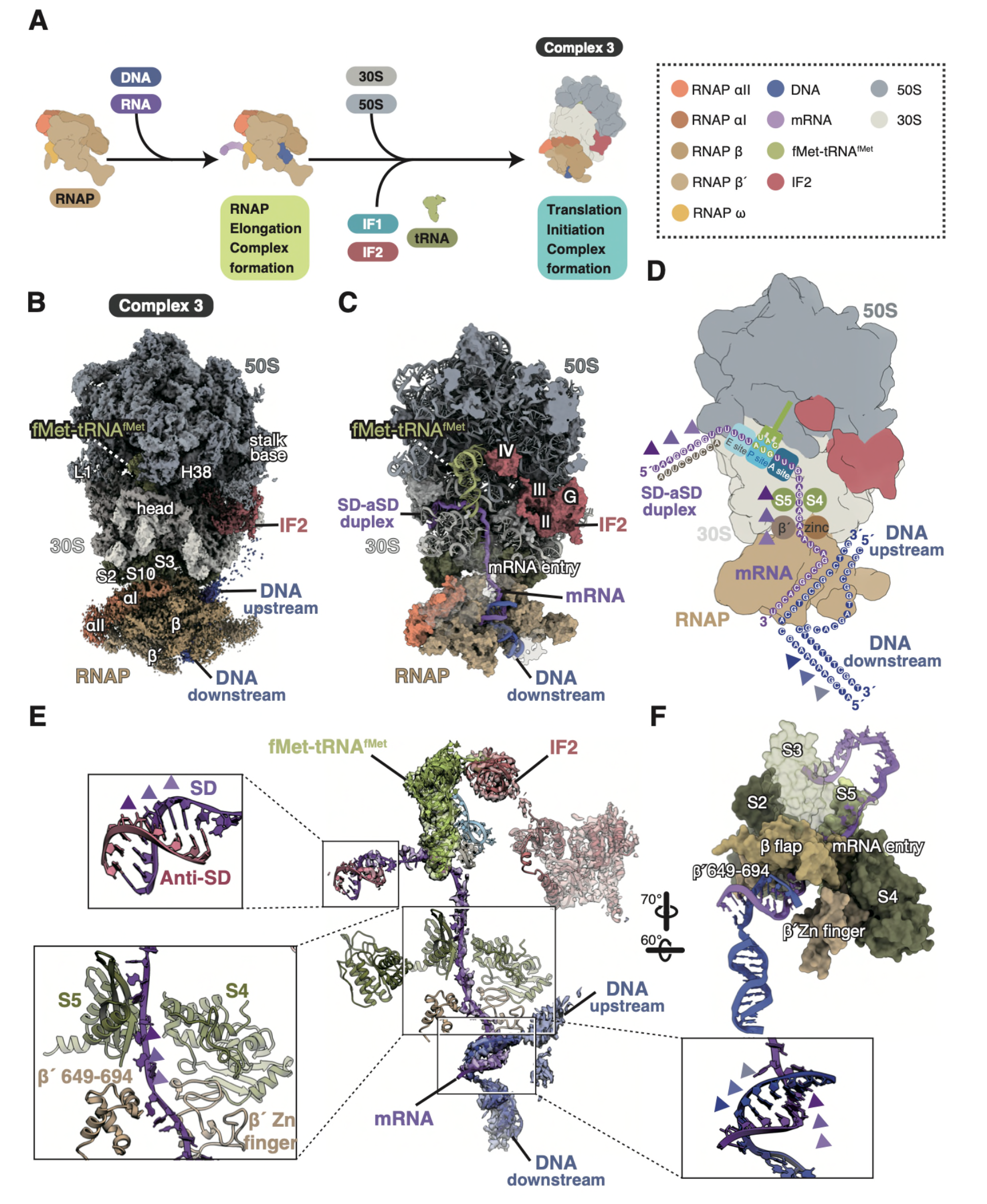
Cryo-EM structure of the 70S initiation complex assembled on RNAP-EC. (**A**) Schematic representation of the assembly of the 70S initiation complex on RNAP-EC in the presence of translational initiation factors for cryo-EM analysis. (**B**) Cryo-EM reconstruction of the 70S initiation complex bound with RNAP-EC to the mRNA entry of the 30S subunit (complex 3) (**C**) Surface representation of the model of complex 3 showing IF2, tRNA and mRNA in this complex with the cross section of the structure. (**D**) Schematic representation of complex 3 showing the DNA/RNA interaction and location inside the complex. (**E**) mRNA inside of the TTC showing the SD duplex formation and the mRNA interactions with the mRNA channel formed by RNAP and the 30S subunit. (**F**) A surface representation of the mRNA channel with mRNA and DNA.

## Discussion

Based on the current structures, we propose a model of the ribosome assembly on RNAP-EC for the establishment of transcription-translation coupling (Fig. 4). First, as transcription proceeds, the 5’-end of the nascent mRNA emerges from the RNA exit channel of RNAP (State I of Fig. 4). The emerged SD sequence immediately forms an RNA duplex with anti-SD within the 30S ribosome subunit, and the duplex is docked into the SD chamber (State II of Fig. 4, corresponding to complex 1). Subsequently, the start codon on the mRNA is clipped by the codon recognition of fMet-tRNA^fMet^, which stabilizes mRNA in the 30S mRNA path. At this point, the ribosomal head and shoulder have an open gap between them, allowing mRNA to pass through the gap to relocate RNAP-EC to the back side of the 30S subunit (State II of Fig. 4). RNAP-EC could undergo a large swinging motion to interact with the “canonical” mRNA entry channel, consisting of the ribosomal proteins S3, S4 and S5. The RNAP αI subunit interaction with the ribosomal proteins S3 and S10 causes a conformational change in the 30S head, which locks the gap closed, placing the mRNA within the canonical mRNA path (State III of Fig. 4, corresponding to complex 2). Alternatively, if RNAP-EC directly binds to the back side of the 30S subunit via the mRNA channel formation, mRNA could be inserted directly into the mRNA path (State II**’** of Fig. 4). This scenario could occur, for example, in the case of genes lacking the SD sequence (non-SD gene). In *E. coli*, the TTC formation in non-SD and long operons are reportedly mediated by RfaH (*39*). Subsequently, with the assistance of IF2, the 50S ribosomal subunit associates with the 30S subunit to form the 70S initiation complex (State IV of Fig. 4, corresponding to complex 3). The GTP hydrolysis facilitates the IF2 dissociation from the 70S, which would complete the TTC.

**Fig. 4.**
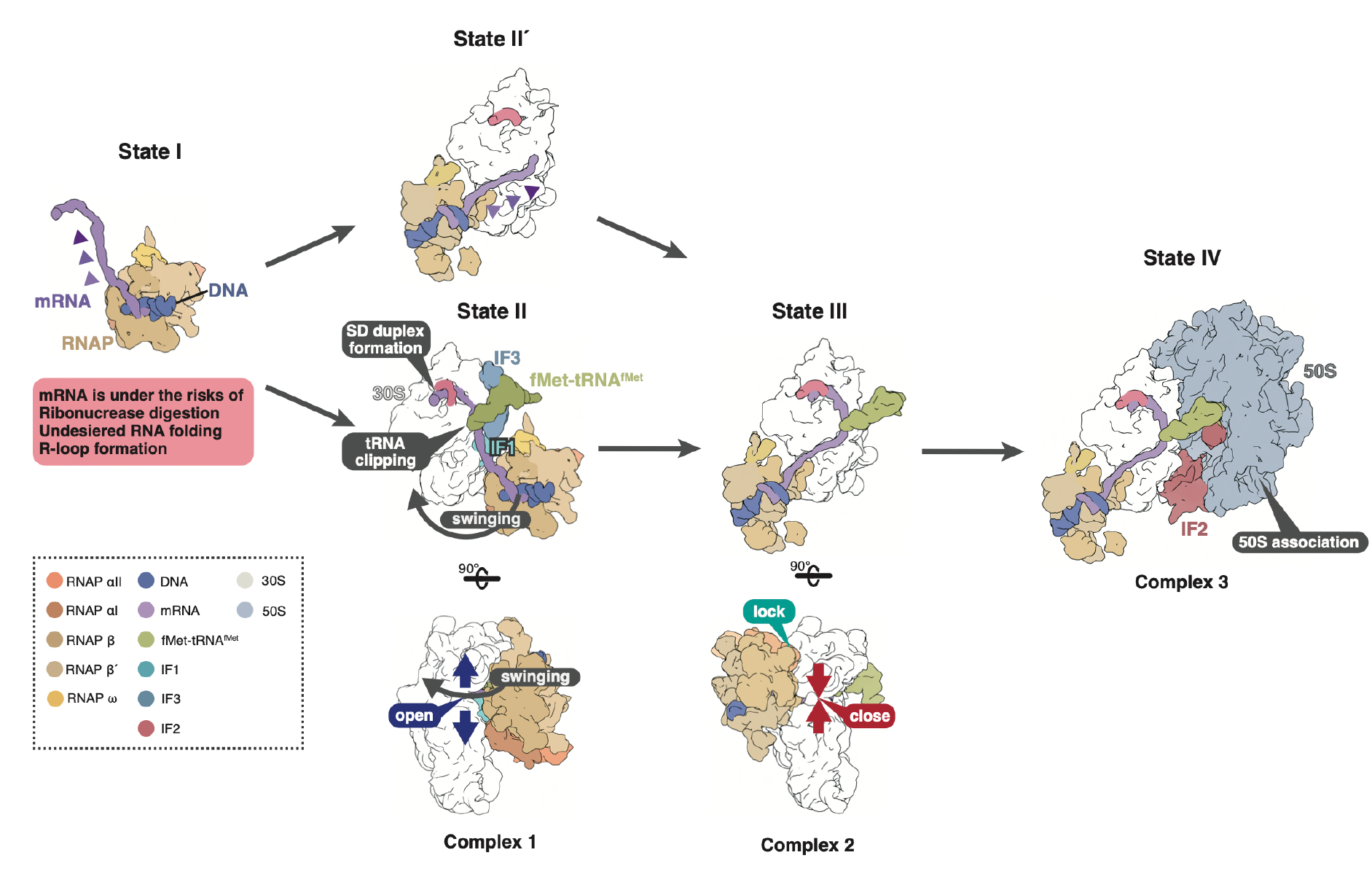
Model of the transcription-translation complex establishment. (State I) the RNAP starts transcription while the nascent mRNA is exposed from the RNA exit channel, increasing the risk of ribonuclease attack, undesired RNA folding, and R-loop formation. (State II) The exposed SD sequence forms the duplex with the antiSD sequence, and the initiator tRNA binding clips the mRNA on the 30S subunit (top). At this point, the mRNA channel is in an open conformation to allow the RNAP-EC to swing (bottom). (State II**’**) The alternative pathway for the non-SD gene. The RNAP could directly interact with the 30S subunit to insert the nascent RNA toward the mRNA path. (State III) The RNAP-EC is relocated and attached to the mRNA channel on the 30S subunit (top). This interaction locks the 30S subunit in a closed conformation (bottom). (State IV) IF2 supports the 50S binding to the RNAP-EC-30S initiation complex to establish the translation elongation competent TTC.

As bacterial mRNAs do not contain the 5’-cap structure, the mRNA 5’-end exiting the RNA-exit channel of RNAP is immediately susceptible to nuclease digestion. The RNAP-30S interactions and direct mRNA transfer between them observed in both complexes 1 and 2 would prevent the access of ribonucleases, and thus, mRNA degradation. This is consistent with the notion that translation initiation protects mRNA from digestion by RNaseJ (*23*)(*24*). The unprotected RNA 5’-end could also undergo secondary structure formation within the mRNA or R-loop formation with DNA, which affects transcription elongation. The direct incorporation of mRNA into the 30S mRNA path, as observed in complexes 1 and 2, could also preclude the possibility of such RNA folding and R-loop formation. This is analogous to the transcription anti-termination mechanism in rRNA biogenesis by the rRNA transcription anti-termination complex (*rrn*TAC), which binds the RNA-exit site of RNAP and serves as an RNA chaperone (*40*).

The RNAP-30S complex formation may also prevent premature termination of transcription mediated by the transcription termination factor Rho. Rho binds a pyrimidine-rich, free region of mRNA, and accesses the RNA-exit site of RNAP-EC to terminate transcription (*41*)(*42*). Uncoupling of RNAP from the ribosome is known to trigger the Rho recruitment to RNAP-EC to induce premature termination (e.g. the polarity effect (*27*)). The immediate mRNA incorporation into 30S (as in both complexes 1 and 2) could block the access by Rho and prevent premature transcription termination.

In this study, we visualized the course of translation initiation complex formation coupled with mRNA transcription by RNAP. The obtained cryo-EM structures represent a seamless mRNA transfer from RNAP to the ribosome, avoiding the mRNA degradation and unnecessary interactions.

## Acknowledgments

The cryo-EM experiments were performed with the Krios G4 microscope at the RIKEN Yokohama Cryo-EM facility. We thank Takashi Osanai (OSA) for setting up workstations for cryo-EM data analysis.

## Author contributions

Conceptualization: TY, SS

Methodology: TY, YM, TU, YT, AN, TS, MS, SS

Investigation: TY, YM, TU, YT, AN

Visualization: TY, SS

Funding acquisition: TY, SS

Supervision: SS

Writing – original draft: TY, YM, SS

Writing – review & editing: TY, YM, SS

## Data and materials availability

All data needed to evaluate the conclusions in the paper are present in the paper and/or the Supplementary Materials. The cryo-EM maps and coordinates have been deposited in the Electron Microscopy Data Bank (EMDB) and Protein Data Bank (PDB), respectively, with the following accession codes: EMD-39123/PDB 8YBU (complex 1), EMD-39124/PDB 8YBV (complex 2), EMD-39125/PDB 8YBW (complex 3).

## Materials and Methods

### Proteins

The RNAP core enzyme was purified from *T. thermophilus* cells, as described previously(*43*). Briefly, the RNAP was partially purified from the cell lysate by polyethyleneimine extraction, and the core enzyme was separated from the holoenzyme by cation-exchange chromatography using SP Sepharose FF (Cytiva). The core enzyme was further purified by heparin affinity, anion-exchange, and size exclusion chromatography steps using HiTrap Heparin HP, Resource Q and Superdex 200 columns (Cytiva).

*T. thermophilus* NusG gene was cloned into pET47b vector. The N-terminally His-tagged NusG was expressed in *E. coli* Rosetta2 (DE3). The cells were disrupted in the lysis buffer containing 50 mM Tris-HCl pH 8.0, 150 mM NaCl and 1 mM DTT. The lysate was heated at 80°C for 10 minutes, and the supernatant was loaded onto a Ni affinity column (Ni-Sepharose 6 FF, Invitrogen). The His-tag was cleaved by the HRV3C protease on the column, and NusG was eluted with the lysis buffer.

*T. thermophilus* translational initiation factors 1, 2 and 3 were expressed and purified as previously described, except that after heat treatment IF2 was purified using the HiTrap Q column and further purified using the resource Q column followed by the Mono Q column(*29*). Plasmids encoding IF1 and IF3 were provided from the biological resource of RIKEN(*44*).

### Initiator tRNA

Formyl-methioninyl (fMet)-tRNA^fMet^ was isolated from the total RNA of *E. coli DRNA* strain obtained from KO collection(*45*) by reciprocal circulating chromatography (RCC) method(*46*). DNA probes (5**’**-GTTATGAGCCCGACGAGCTACCAGGCTGCT-3**’**) with a 5**’**-ethylcarbamate amino linker (Sigma-Aldrich) were covalently immobilized on NHS-activated Sepharose 4 Fast Flow (GE Healthcare) and packed into tip columns for the RCC instrument. To suppress the deacylation of fMet moiety from the 3**’**-terminus of tRNA^fMet^ and the oxidation of methionine the RCC method was performed in the binding buffer containing 1.2 M NaCl, 30 mM MES-NaOH (pH 6.0), 15 mM EDTA and 2.5 mM DTT by pipetting 17 times in the RNA binding process. To analyze of 3**’**-terminus RNA fragment from the fMet-tRNA^fMet^, the isolated tRNA^fMet^ was digested with RNase T_1_ and analyzed by capillary LC/nanoESI MS system.

### *T. thermophilus* ribosomes

*T. thermophilus* cells in the pellet were disrupted by freeze-thawing in RBS buffer containing 20 mM HEPES-KOH pH 7.6, 10 mM Mg(OAc)_2_, 30 mM NH_4_Cl, and 250 µg/ml Lysozyme. Cell debris was cleared by centrifugation for 20 mins at 20,000 x *g*. Subsequently, crude ribosomes in the supernatant were pelleted by ultracentrifugation for 2 hours at 134,000 x *g*. The pellet was resuspended with RBS buffer containing lower magnification (0.5 mM) for the dissociation of ribosomal subunits. Ribosomal subunits were pelleted by ultracentrifugation for 4 hours at 134,000 x *g*. The pellet was resuspended with RBS buffer (Mg 0.5 mM) and applied for 10% to 40% (w/v) sucrose density gradient centrifugation. Ultracentrifugation was performed for 16 hours at 20,000 rpm using a SW28 centrifuge swing rotor (Beckman Coulter). The 30S ribosome and 50S ribosome were fractionated and pelleted by ultracentrifugation for 4 hours at 134,000 x *g*. Pellets were resuspended in the RBS buffer for subsequent processes.

### Reconstitution of the RNAP elongation complex for cryo-EM

DNA sequences used for the RNAP-EC formation for 30S initiation complexes are as follows: the template DNA: 5**’**-CTGAAGATCGAAAAAAGCACGTGCGGCCTCGCGTGGTGTAGGAGCTAAGCGTCTGGATACCGAGCATTGC-3**’**, the non-template DNA: 5**’**-GCAATGCTCGGTATCCAGACGCTTAGCTCCTACACCACGCGGGCGGTAGCACGCTTTTTTCGATCTTCAG-3**’**. DNA sequences used for the RNAP-EC formation for 70S initiation complexes are as follows: the template DNA: 5**’**-GAAGATCGAAAAAAGCACGTGCGGCCtCGCGTGGTGTAGGAGCTAAGC-3**’**, the non-template DNA: 5**’**-GCTTAGCTCCTACACCACGCGGGCGGTAGCACGCTTTTTTCGATCTTC-3**’**. The RNA sequence for the RNAP-EC formation for the 30S and 70S initiation complexes is the 43 nt mRNA: 5**’**-UAAGGAGGUUUUUUAUGUUUGUAGUAGAAAUCAGGCCGCACGU-3**’**

For reconstitution of the RNAP elongation complex, the DNA/RNA scaffold was annealed by incubating the mixture of template DNA, non-template DNA, and RNA at 80°C for 2 minutes and gradually cooling to 25°C. 400 nM RNAP and 500 nM DNA/RNA scaffold were mixed in 100 µl of buffer containing 20 mM HEPES-NaOH pH 7.5 and 100 mM NaCl. After the mixture was incubated at 25 °C for 5 minutes, excess DNA and RNA were removed by size exclusion chromatography using a Superose6 Increase column (Cytiva).

### Complexes formation for cryo-electron microscopy

1) Complex formation of the RNAP-EC bound 30S initiation complex.

30S ribosome, RNAP-EC, IF1, IF3, fMet-tRNA^fMet^ and NusG were mixed in the buffer containing 20 mM HEPES-KOH pH 7.6, 10mM Mg(OAc)_2_, 30mM NH_4_Cl and incubated at 37°C for 10 min at the final concentration of 100 nM, 1 µM, 1µM, 1µM, 300 nM, 1 µM respectively. Subsequently, IF2 and GDPNP at the final concentration of 1 µM and 200 µM were added and further incubated at 37°C for 10 min.

2) Complex formation of the RNAP-EC bound 70S initiation complex.

30S and 50S ribosome, RNAP-EC, fMet-tRNA^fMet^, and NusG were mixed in the buffer containing 20 mM HEPES-KOH pH 7.6, 10mM Mg(OAc)_2_, 30mM NH_4_Cl and incubated at 37°C for 30 min at the final concentration of 50 nM, 100 nM, 120 nM, 100 nM, respectively. Subsequently, IF1, IF2, and GDPNP at the final concentration of 500 nM, 500 nM, and 200 µM were added and further incubated at 37°C for 5 min.

### Cryo-EM grid preparation, data acquisition and image processing

Quantifoil R1.2/1.3 300 mesh Cu grids (Quantifoil Micro Tools GmbH) were coated with a homemade amorphous carbon film, which is prepared by vacuum deposition using a JEE-420 vacuum evaporator (JEOL). These carbon-coated grids were glow discharged at 5 mA (soft condition) for 10 s with a PIB-10 Plasma Ion Bombarder (Vacuum Device). A 3 µL portion of RNAP-ribosome complexes at the concentration of 50 nM were applied onto the carbon-coated surface of grids, and they underwent vitrification by Vitrobot Mark IV (Thermo Fisher Scientific) plunge-freezing device.

1) The RNAP-EC bound 30S initiation complex.

The automated data acquisition was performed with EPU program controls a Krios G4 transmission electron microscope (Thermo Fisher Scientific) equipped with a BioQuantum energy filter and K3 camera direct electron detector (Gatan). The microscope was operated at 300 kV accelerating voltage. The total exposure dose at the specimen was 47 e^-^Å^-2^ and recorded on 47 continuous frames. Motion correction was performed with the motion correction program implemented in Relion 3.0(*47*). The CTF parameters of motion corrected micrographs were estimated with the CFTFIND4 program. 3,147K particles were picked from 24,970 motion-corrected micrographs (fig. S2). This dataset was subsequently performed with 2D classification and converged into 2D class-averaged images showing 30S complexes (2,165K particles were selected). 3D classification was performed, and the dataset was sorted into five major sub-groups. Three classes were comprised of 177K (branch 1), 224K (branch 2), and 208K particles (branch 3), respectively, showing RNAP binding on the 30S ribosomal subunit with translation initiation factors. 82K particles from branches 2 and 3 with bindings of RNAP, IF1, IF3, and tRNA were further refined, following CTF-refinement, Bayesian Polishing, and Auto-refinement. The focused classification on the RNAP density sorted 40K particles with the RNAP binding on the 30S inter-subunit side. Another round of focused classification sorted 4.4K particles having stable RNAP. This class was refined with original unsubtracted particles, and the reconstructed complex 1 structure at the resolution of 6.7 Å. Local-resolution estimation was performed with ResMap program(*48*) and shown in panel C. 69K particles having the RNAP density at the mRNA entry side (branch 5) was refined. Subsequent 3D classification sorted the particles with the stable RNAP binding (54K particles). This class followed Auto-refinement, CTF-refinement, Bayesian Polishing, and another round of Auto-refinement reconstructed complex 2 structure at the resolution of 3.6 Å. Local-resolution estimation was performed with ResMap program(*48*) and shown in panel E.

2) The RNAP-EC bound 70S initiation complex.

The automated data acquisition was performed with the SerialEM program(*49*), which controls a Tecnai Arctica transmission electron microscope (Thermo Fisher Scientific) equipped with a K2 summit direct electron detector (Gatan). The microscope was operated at 200 kV accelerating voltage. The total exposure dose at the specimen was 50 e^-^Å^-2^ and recorded on 40 continuous frames. Motion correction was performed with the motion correction program implemented in Relion 3.0. The CTF parameters of motion-corrected micrographs were estimated with the CFTFIND4 program. Particle picking of ribosome particles was performed with the Gautomatch program. Subsequent image processing was performed with Relion 3.0 package(*47*). 1,699K particles were picked from 8,148 motion-corrected micrographs (fig. S7). This dataset was divided into 7 groups, and each sub-group was image-processed independently. After 2D classification, consensus reconstruction and 3D classification were performed on each dataset. Particles showing a tRNA, weak RNAP, and weak IF2 densities were joined and further performed with auto-refinement (fig. S7). This dataset was 3D classified into 4 groups. One class consisting of 36K particles having strong densities for RNAP, IF2, and tRNA was performed with CTF-refinement, Bayesian Polishing and auto-refinement to obtain the cryo-EM structure at 3.7Å. Local-resolution estimation was performed with ResMap program(*48*) and shown in panel C.

### Model building and figure preparation

Structural models were constructed using the COOT and ISOLDE programs based on previously reported structures of *T. thermophilus* 70S ribosome (PDB ID: 4V4Y)(*50*), RNAP-DNA-RNA (PDB ID: 4WQS)(*17*), IF1, IF2, IF3, fMet-tRNA^fMet^ (PDB ID: 5LMV)(*29*) . The topology of the models were refined into cryo-EM maps using phenix.real_space_refine(*51*). Figures showing cryo-EM densities and constructed atomic models were depicted by using either UCSF Chimera(*52*) or UCSF ChimeraX(*53*).

