## Supplementary materials for "Structural insight into bacterial co-transcriptional translation initiation"

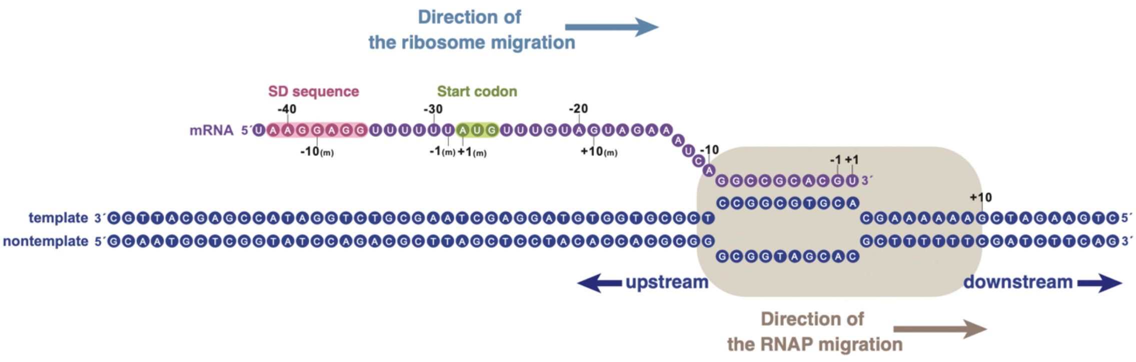


**Fig. S1.**

**Sequences of the RNA and DNAs in the RNAP-EC used for complex formation of the 30S initiation complex.** A 43 nt RNA is shown in purple with an SD sequence and a start codon. Nucleotides +1 to -9 form Watson-Click base pars with the template DNA. The template DNA from nucleotides from -51 to -10 (upstream) and from +2 to +19 (downstream) form base pars with the nontemplate DNA on the RNAP. Direction of the ribosome and the RNAP migrations are shown as arrows.

A

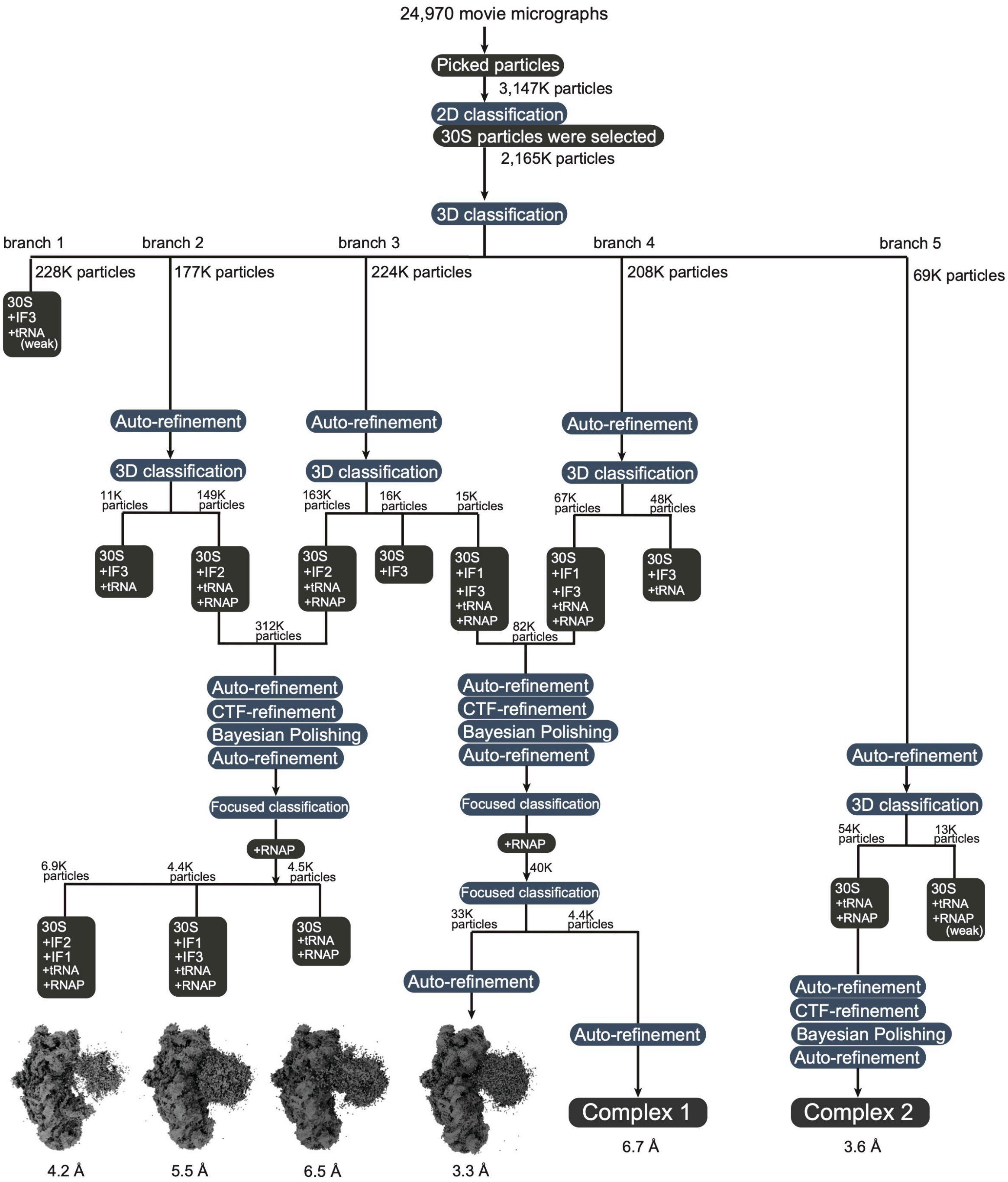
24,970 movie micrographs
Picked particles
3,147K particles
2D classification
30S particles were selected
2,165K particles
3D classification
branch 1
branch 2
branch 3
branch 4
branch 5
228K particles
177K particles
224K particles
208K particles
69K particles
30S +IF3 +tRNA (weak)
Auto-refinement
3D classification
11K particles
149K particles
163K particles
16K particles
15K particles
67K particles
48K particles
30S +IF3 +tRNA
30S +IF2 +tRNA +RNAP
30S +IF2 +tRNA +RNAP
30S +IF3
30S +IF1 +IF3 +tRNA +RNAP
30S +IF1 +IF3 +tRNA +RNAP
30S +IF3 +tRNA
312K particles
82K particles
Auto-refinement
CTF-refinement
Bayesian Polishing
Auto-refinement
Focused classification
+RNAP
6.9K particles
4.4K particles
4.5K particles
30S +IF2 +IF1 +tRNA +RNAP
30S +IF1 +IF3 +tRNA +RNAP
30S +tRNA +RNAP
40K
33K particles
4.4K particles
Complex 1
6.7 Å
54K particles
13K particles
30S +tRNA +RNAP
30S +tRNA +RNAP (weak)
Complex 2
3.6 Å
4.2 Å
5.5 Å
6.5 Å
3.3 Å

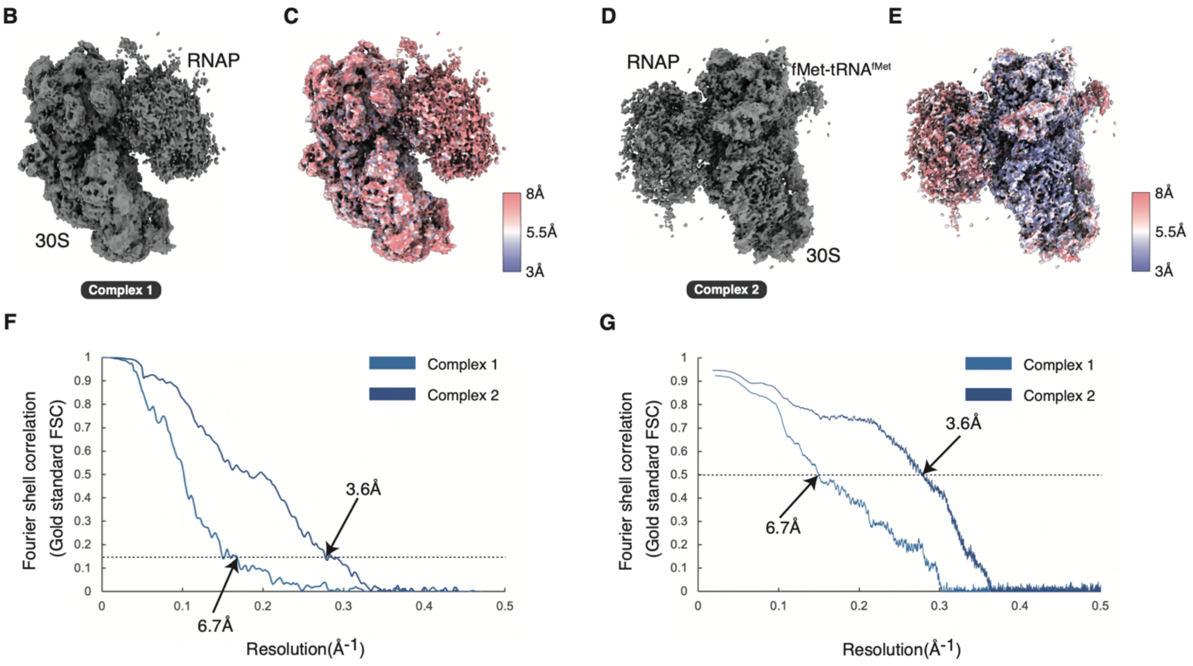


**Fig. S2.**

**Single particle cryo-EM of the RNAP-EC bound to the 30S initiation complex.**

(**A**)The scheme of the image processing of this data set. From 24,970 motion-corrected movie micrographs, 3,147K particles were picked and 2D classification was performed. 30S particles and RNAP-bound particles were selected for the first round of 3D classification, which sorted particles into five major groups. The three classes having 30S ribosomes with the RNAP-EC binding on the inter subunit side were further processed. The particles with IF2 bindings were combined for the refinement. Reconstructed structure having non converged the RNAP density, was subjected to the focused classification on the RNAP. The particles were classified into three major classes with the combination of initiation factors. The particles with IF1 and IF3 bindings were combined for the refinement. This class was focused classified by the density of the RNAP. The subsequent refinement from 4.4K particles reconstructed the complex 1 structure at the resolution of 6.7Å. One of five classes having the RNAP density on the mRNA entry side was refined. The focused classification on the RNAP density was performed to obtain the RNAP-EC-30S structure. The subsequent refinement from 54K particles reconstructed the complex 2 structure at the resolution of 3.6Å. (**B**) cryo-EM map of complex 1. (**C**) Local resolution distribution of the cryo-EM map of panel B. (**D**) cryo-EM map of complex 2. (**E**) Local resolution distribution of the cryo-EM map of panel D. FSC of the cryo-EM reconstruction and map vs model are shown in panels (**F**) and (**G**).

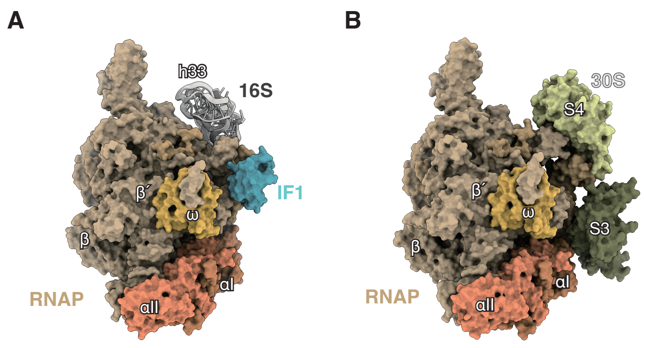


**Fig. S3.**

**The RNAP interactions with the ribosomal components in complex 1 and complex 2.** (**A**) The RNAP interaction in complex 1 with helix 33 of the 16S rRNA and IF1 on the A site. (**B**) The RNAP interaction in complex 2 with ribosomal proteins S3 and S4 forming the mRNA entry channel.

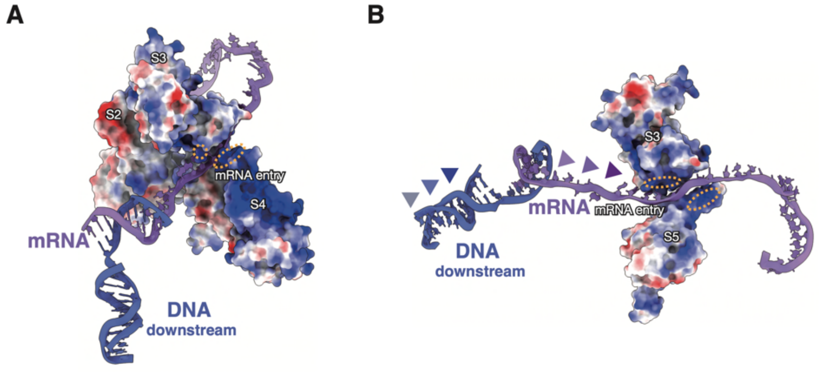


**Fig. S4.**

**Positively charged mRNA path connecting RNAP to 30S.** (**A**) Surface charge distribution of the docking triangle on 30S. Positively charged residues are clustered on the entry of mRNA (dotted orange circle). (**B**) Positively charged residues are also situated inside the portal guiding mRNA insertion into the mRNA path.

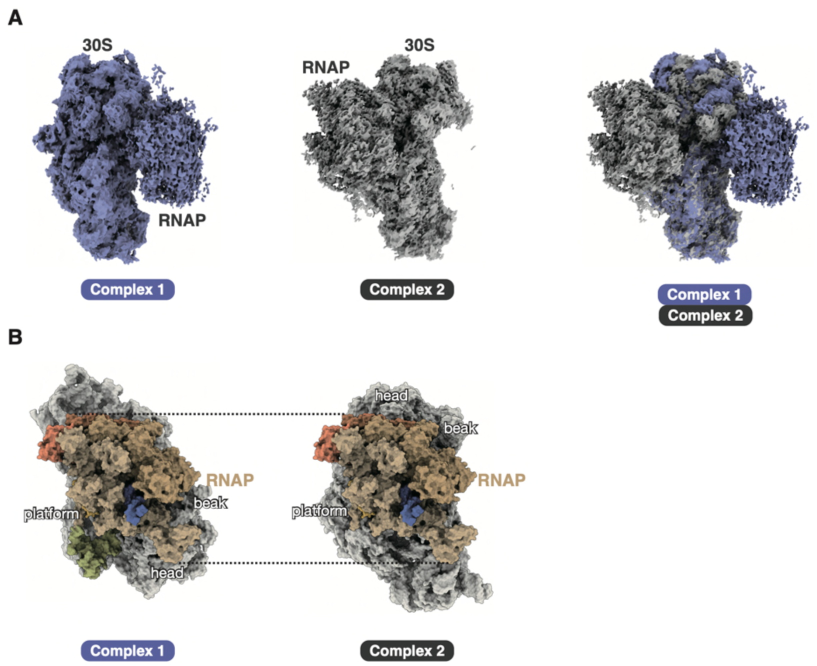


**Fig. S5.**

**The comparison of complexes 1 and 2.** (**A**) cryo-EM maps of complex 1 (left) and complex 2 (middle). These two structures were aligned of the density of the 30S body as a guide. The aligned maps were superimposed and shown for the comparison (right). (**B**) The atomic models of complexes 1 and 2 were aligned by the RNAP portion as a guide. The relative orientation of the 30S with respect to the RNAP is shown. The dotted lines are the guides of the top and bottom edges of the RNAP in this orientation.

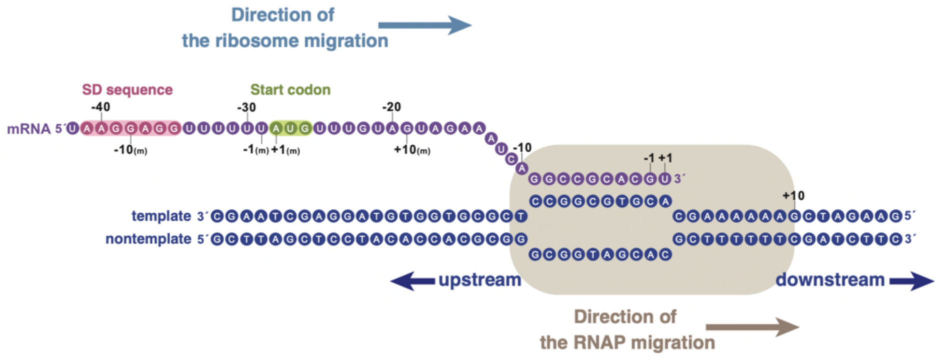


**Fig. S6.**

**Sequences of the RNA and DNAs in the RNAP-EC used for complex formation of the 70S initiation complex.** A 43 nt RNA is shown in purple with an SD sequence and a start codon. Nucleotides +1 to -9 form Watson-Click base pars with the template DNA. The template DNA from nucleotides from -31 to -10 (upstream) and from +2 to +17 (downstream) form base pars with the nontemplate DNA on the RNAP. Direction of the ribosome and the RNAP migrations are shown as arrows.

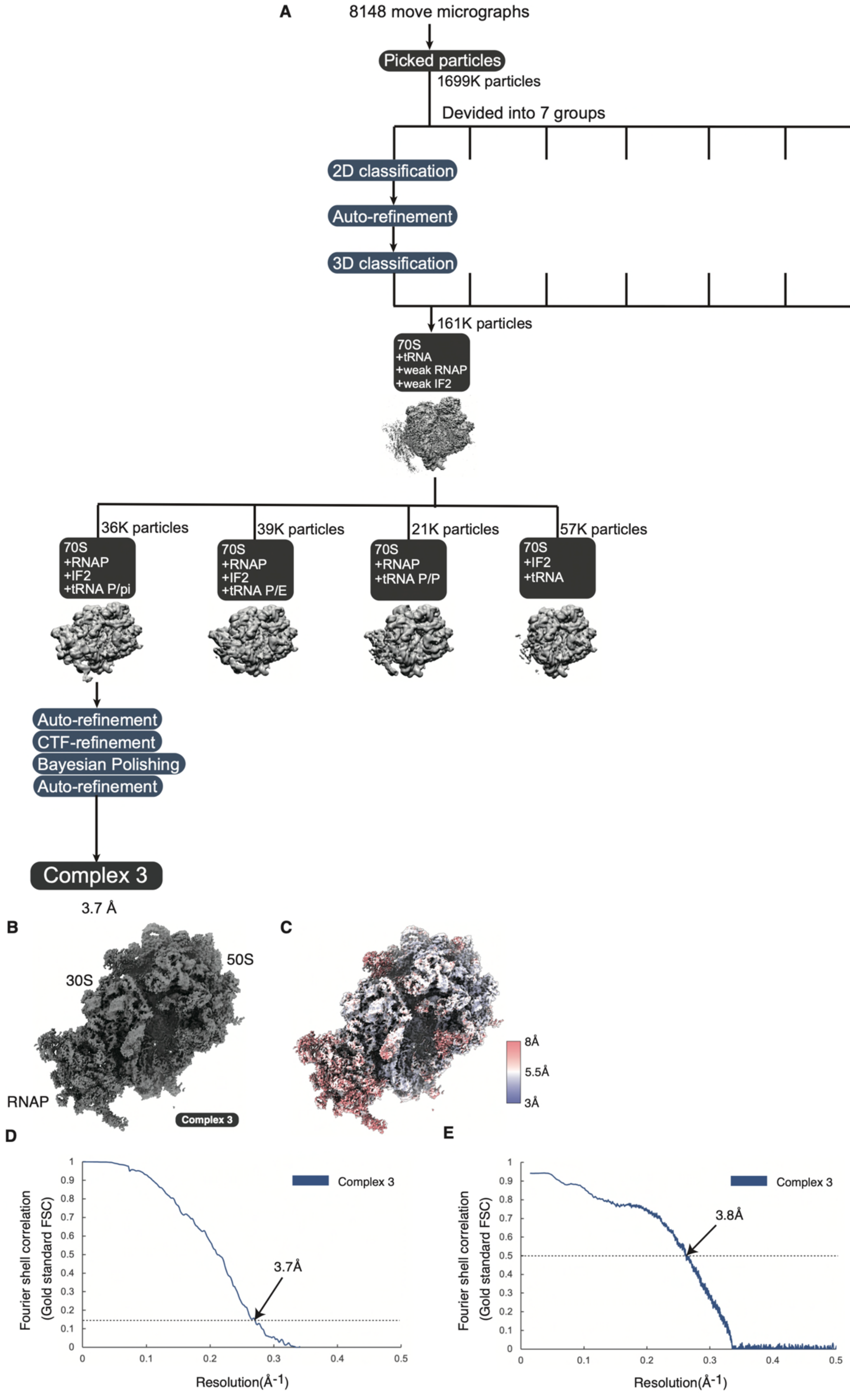

A
8148 move micrographs
Picked particles
1699K particles
Devided into 7 groups
2D classification
Auto-refinement
3D classification
161K particles
70S
+tRNA
+weak RNAP
+weak IF2
36K particles
70S
+RNAP
+IF2
+tRNA P/pi
39K particles
70S
+RNAP
+IF2
+tRNA P/E
21K particles
70S
+RNAP
+tRNA P/P
57K particles
70S
+IF2
+tRNA
Auto-refinement
CTF-refinement
Bayesian Polishing
Auto-refinement
Complex 3
3.7 Å
B
50S
30S
RNAP
Complex 3
C
8Å
5.5Å
3Å
D
Complex 3
3.7Å
Fourier shell correlation
(Gold standard FSC)
Resolution(Å-1)
E
Complex 3
3.8Å
Fourier shell correlation
(Gold standard FSC)
Resolution(Å-1)

**Fig. S7.**

**Single particle cryo-EM of RNAP-EC-bound 70S initiation complex.** (**A**) Scheme of image processing of complex 3. 1,699K particles from 8,148 motion corrected movie micrographs were divided into seven groups and processed in parallel for the matter of the time shortening. Each group underwent 2D classification, auto-refinement and 3D classification to sort particles with RNAP densities. RNAP-bound 70S particles were joined (161K particles) and auto-refined. This cryo-EM structure of the 70S ribosome showed densities for tRNA, weak RNAP and weak IF2. To further classify this sub-population, another round of 3D classification was performed. One of major group consists of the 70S with RNAP, IF2 and tRNA (36K particles). This group underwent CTF-refinement, Bayesian polishing and auto-refinement and reached at the average resolution of 3.7 Å. (**B**) cryo-EM map of complex 3. (**C**) Local resolution distribution of the cryo-EM map of complex 3. FSC of the cryo-EM reconstruction and map vs model are shown in panels (**D**) and (**E**).

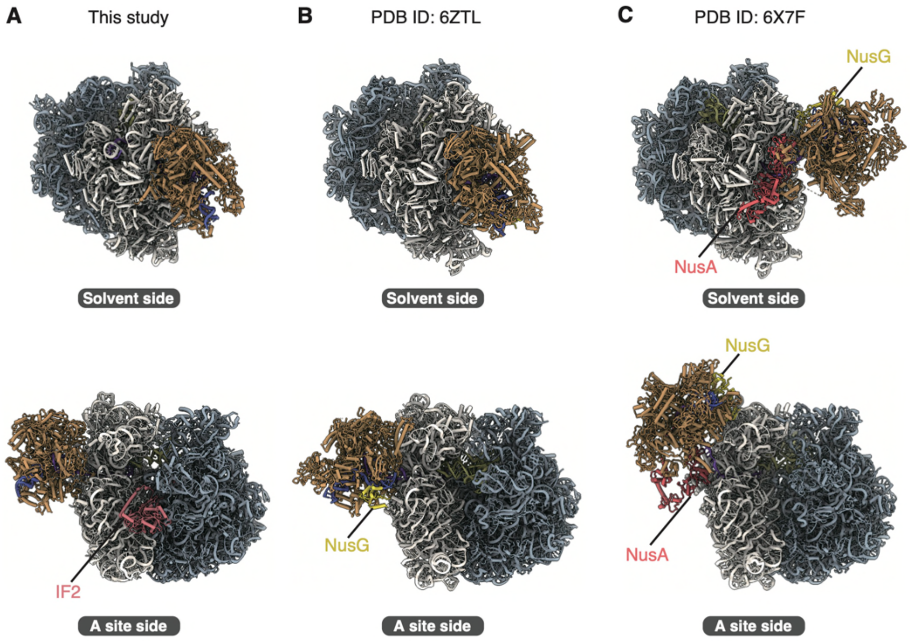


**Fig. S8.**

**The structural comparison of complex 3 with the previously reported TTC structures.** (**A**) The structure of the RNAP-EC bound 70S initiation complex (complex 3, this study). (**B**) The structure of the RNAP collided state connected to the 70S via NusG.(*10*) (**C**) The structure of the RNAP in coupled state connected to the 70S via NusA and NusG(*9*).

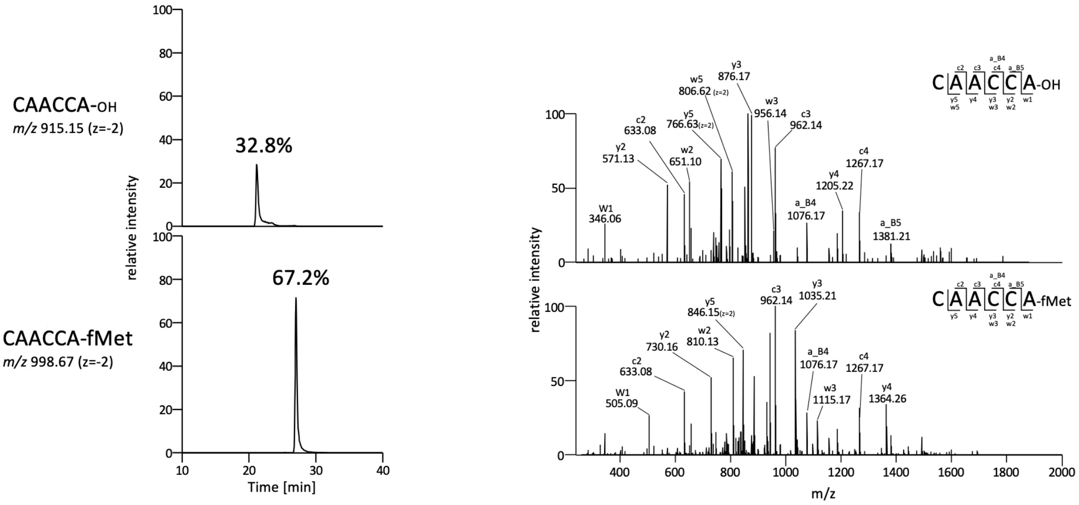


**Fig. S9.**

**RNA-MS analysis of fMet-tRNA$^{fMet}$ isolated by RCC method.** Extracted ion chromatograms (XIC) (left panels) and CID spectrum (right panels) of the 3′-terminus fragments of deacylated tRNA$^{fMet}$ (upper panel) and fMet-tRNA$^{fMet}$ (lower panel) digested with RNase $T_1$. Sequence, m/z value, and charge state of each fragment are indicated in each panel. The abundance ratio of fMet-tRNA$^{fMet}$ was calculated from the ratio of the XIC peak areas of the two fragments.

**Table S1.**

Cryo-EM data collection, image processing, model building and validation statistics.

| | (Complex 1)<br>(EMDB-39123)<br>(PDB 8YBU) | (Complex 2)<br>(EMDB-39124)<br>(PDB 8YBV) | (Complex 3)<br>(EMDB-39125)<br>(PDB 8YBW) |
|---|---|---|---|
| **Data collection and processing** | | | |
| Magnification | 105,000 | | 23,500 |
| Voltage (kV) | 300 | | 200 |
| Electron exposure ($e^{-}$/Å$^2$) | 47 | | 50 |
| Defocus range (μm) | 0.5-3.5 | | 0.5-3.5 |
| Pixel size (Å) | 0.83 | | 1.47 |
| Symmetry | $C_1$ | | |
| Initial particle images (no.) | 3,147,199 | | 1,699,349 |
| Final particle images (no.) | 4,425 | 53,806 | 35,959 |
| Map resolution (Å) | 6.7 | 3.6 | 3.6 |
| FSC threshold | 0.143 | 0.143 | 0.143 |
| Map sharpening *B* factor (Å$^2$) | -10 | -10 | -40 |
| **Refinement** | | | |
| Model composition | | | |
| Non-hydrogen atoms | 80,149 | 78,079 | 173,244 |
| Protein residues | 5,566 | 5,306 | 9,066 |
| RNA residues | 1,669 | 1,669 | 4,703 |
| Mg | 63 | 77 | 300 |
| Zn | 2 | 2 | 2 |
| R.m.s. deviations | | | |
| Bond lengths (Å) | 0.003 | 0.004 | 0.006 |
| Bond angles (°) | 0.671 | 0.641 | 0.850 |
| Validation | | | |
| MolProbity score | 2.79 | 2.73 | 3.11 |
| Clashscore | 62.99 | 52.81 | 55.27 |
| Poor rotamers (%) | 0.32 | 1.05 | 2.57 |
| Ramachandran plot | | | |
| Favored (%) | 90.98 | 91.16 | 88.74 |
| Allowed (%) | 8.96 | 8.84 | 11.05 |
| Disallowed (%) | 0.05 | 0.00 | 0.20 |
